# Synaptonemal Complex dimerization regulates chromosome alignment and crossover patterning in meiosis

**DOI:** 10.1101/2020.09.24.310540

**Authors:** Spencer G. Gordon, Lisa E. Kursel, Kewei Xu, Ofer Rog

## Abstract

During sexual reproduction the parental homologous chromosomes find each other (pair) and align along their lengths by integrating local sequence homology with large-scale contiguity, thereby allowing for precise exchange of genetic information. The Synaptonemal Complex (SC) is a conserved zipper-like structure that assembles between the homologous chromosomes. This phase-separated interface brings chromosomes together and regulates exchanges between them. However, the molecular mechanisms by which the SC carries out these functions remain poorly understood. Here we isolated and characterized two mutations in the dimerization interface in the middle of the SC zipper in *C. elegans*. The mutations perturb both chromosome alignment and the regulation of genetic exchanges. Underlying the chromosome-scale phenotypes are distinct alterations to the way SC subunits interact with one another. We propose that the SC brings homologous chromosomes together through two biophysical activities: obligate dimerization that prevents assembly on unpaired chromosomes; and a tendency to phase-separate that extends pairing interactions along the entire length of the chromosomes.

## Introduction

In most cells, homologous chromosomes (homologs) do not physically interact. However, in meiotic cells, homologs are regulated precisely and synchronously as a pair via direct physical interaction, enabling them to exchange genetic information. For these interactions to occur, meiotic chromosomes undergo large scale conformational changes. First, each homolog becomes elongated and assembles onto a stiff proteinaceous structure called the axis. Then, homologs pair and align from end to end. Finally, homologs exchange genetic information through generation of crossovers. Crossovers also provide the physical linkages that hold the homologs together and allow them to segregate correctly into the gametes. Therefore, failure to bring the homologs together, and the subsequent absence of crossovers, result in aneuploid gametes and sterility.

Homolog alignment and regulation of crossovers rely on the assembly of a specialized interface called the Synaptonemal Complex (SC). (Throughout, we refer to the protein interface assembling between the axes, which is classically defined as the Central Region of the SC, simply as ‘the SC’). The SC loads at sites of localized interactions and processively zips up and aligns homologs along their entire length [1–3]. In addition to aligning homologs, the SC is required for both crossover formation, through recruitment of pro-crossover factors, and for regulating crossover number and distribution [4–6]. One manifestation of this regulation is wide spacing of crossovers through ‘crossover interference’. In *C. elegans*, interference ensures that, regardless of the number of DNA breaks, only one break per chromosome is repaired as a crossover [4]. The mechanism by which the SC implements crossover interference remains unknown.

The SC ladder-like ultrastructure, as it appears in electron micrographs, is conserved across almost all eukaryotes with the two parallel axes separated by 100-150nm (Fig. 1A; [3]). The SC is composed of a handful of subunits (six in *C. elegans* [7–12]) that occupy stereotypic locations relative to the axes (Fig. 1A; [12–16]). However, we have limited understanding of how SC proteins interact with one another, or how their structure helps them to carry out functions such as aligning the homologs or regulating crossovers. Two main challenges have hampered progress. First, although SC ultrastructure and SC functions are conserved, the primary sequences of SC proteins diverge beyond recognition across eukaryotic lineages [3]. Second, attempts to purify SC components or express them *in vitro* have been mostly unsuccessful, and we therefore lack extensive structural and biochemical characterization of the SC [17]. Underlying the latter challenge might be the physical features of the SC: despite its appearance as an ordered, crystalline interface, the SC has recently been shown to have liquid-like properties and mobile subunits [18, 19].

**Figure 1.**
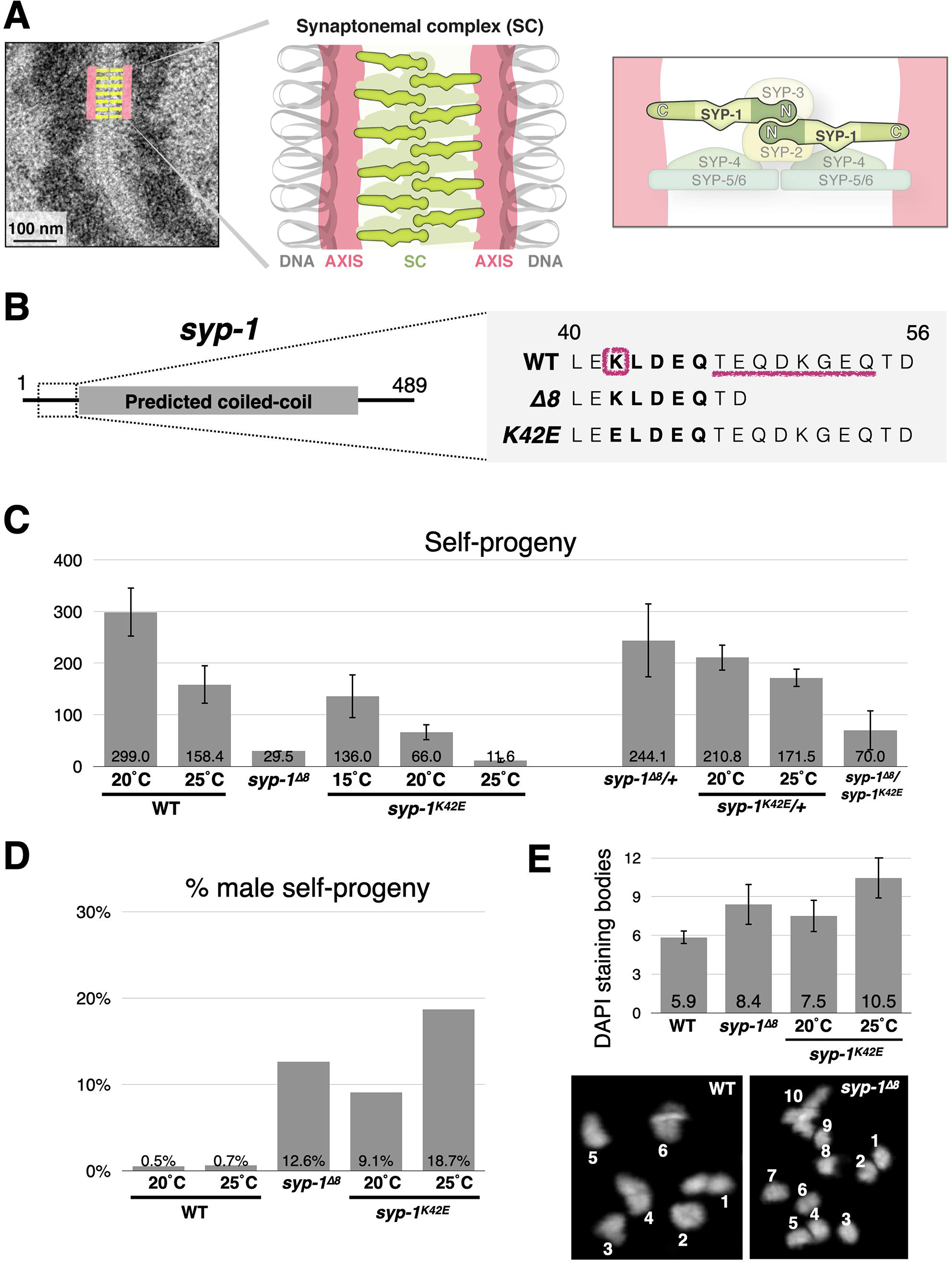
Isolation of separation-of-function mutations in the N-terminus of *syp-1*. A) Model of the SC. Left, electron micrograph of *C. elegans* SC (adapted from [18]) with an overlaid interpretive diagram. The electron dense mass to the sides of the SC is chromatin. Middle, cartoon drawing of the SC. Parallel axes (salmon) organize the chromatin (gray) of each homolog into an elongated structure. The SC (green) holds the two homologs in alignment. Right, the stereotypic organization of SC components (from [12, 13, 15, 16]). The right and left sides of the SC are mirror image of each other. The N- and C-termini of SYP-1 are labeled. B) *syp-1* gene model and mutations. The predicted coiled-coil in the middle is denoted by a gray bar. The conserved 5 amino acid stretch (KLEDQ at positions 42-45) is shown in bold. The residues mutated in *syp-1^K42E^* and *syp-1*^Δ*8*^ are denoted with pink lines. See Fig. S1A-C for more details. C) *syp-1* mutants show fertility defects. Self-progeny counts of wildtype and mutant hermaphrodites. Error bars indicate standard deviation. D) *syp-1* mutants show fertility defects. Percent males among the self-progeny of wildtype and mutant hermaphrodites. Error bars indicate standard deviation.s E) Chromosomes (“DAPI staining bodies”) at the diakinesis stage, immediately preceding the first meiotic division, are more numerous in *syp-1* mutants compared with wildtype, indicating failure at forming crossovers is some chromosomes. Average DAPI staining body counts (reflecting the number of paired and unpaired chromosomes) from the indicated genotypes are shown. Error bars indicate standard deviation. N>14 nuclei for each condition. All pairwise comparisons are significant (Student’s t-test, p<0.05). Bottom, representative images of diakinesis nuclei with counted DAPI staining bodies from wildtype and *syp-1*^Δ*8*^ animals.

Null mutants of single SC components have not been very informative for elucidating structure-function relationships because they commonly abolish the SC altogether. Genetic analyses in multiple model systems, however, have shown that the width of the SC is determined, at least in part, by the length of predicted coiled-coils in SC subunits referred to as transverse filament proteins [20–22], and that the C-termini of these proteins are required for axis associations [23, 24]. In worms these transverse filament proteins include SYP-1 and SYP-5/6, which span, in a head-to-head orientation, the ~100 nm between the axes [7, 12, 16]. The contributions of other domains to specific functions or structural features are much less clear. For example, the ability of the SC to hold paired axes together was suggested to involve, based on *in vitro* analysis, dimerization and cooperative assembly of the N-terminus of the mammalian transverse filament protein SYCP1 [17], and based on genetic data, the N-terminus of budding yeast Zip1p [25, 26]. However, while several genetic perturbations in *C. elegans* cause the SC to be associated with all chromosomes without holding paired axes together [10, 27, 28], the molecular details underlying pairing interactions have not been elucidated. Likewise, despite the identification of conditions that decrease crossover interference - mutations in the C-termini of *syp*-*4*, -*5* and -*6*, and partial depletion of SC components [4, 11, 13] - the molecular mechanisms that generate this chromosome-scale phenotype remain unknown.

To gain mechanistic insight into SC functions, we randomly mutagenized an N-terminal stretch of the SC component SYP-1, which localizes to the middle of the SC (Fig. 1A; [5, 13, 15]). Detailed analyses of the separation-of-function mutations that we isolated showed that the N-terminus of SYP-1 is critical for the coupling of SC assembly and chromosome alignment, as well as for the regulation of crossovers. We propose a model for how SC subunit interactions regulate homolog pairing and genetic exchanges.

## Results and Discussion

### Generating SC hypomorphs by mutating the conserved N-terminus of SYP-1

Alignment of SYP-1 sequences from 17 *Caenorhabditis* species revealed that, despite the low overall sequence conservation, some regions are conserved. These include a phosphorylation site near the C-terminus [29, 30] and sequences required for N-acetylation ([31] and Fig. S1A-B). We focused on a conserved five amino acid stretch at the N-terminus of SYP-1 near the beginning of the predicted coiled-coil (Fig. 1B). We generated mutations by randomly substituting single amino acids using CRISPR/Cas9, giving rise to 50 heterozygous mutants. We then attempted to homozygose these mutants, and eliminated those that were rendered sterile, assuming them to be *syp-1* null mutations [7]. We sequenced the remaining mutants and analyzed them for strong meiotic defects (Fig. 1B and Fig. S1C). Interestingly, many non-conservative substitutions did not exhibit obvious phenotypes. However, two mutants stood out: a lysine to glutamic acid substitution at position 42 (*syp-1^K42E^*), and a serendipitous eight amino acid deletion immediately downstream of the conserved stretch we targeted for mutagenesis (*syp-1*^*Δ8*^; Fig. 1B).

Both mutants exhibited hallmarks of meiotic defects. Self-progeny viability was dramatically reduced, and was accompanied by a high ratio of male progeny indicative of X chromosome missegregation (Fig. 1C-D). The number of chromosomes prior to the meiotic divisions (‘DAPI staining bodies’) was larger than the six linked pairs observed in wildtype animals (Fig. 1E). Notably, we found *syp-1^K42E^* to be temperature sensitive. Progeny of *syp-1^K42E^* animals exhibited progressively decreased viability and increased incidence of males with increasing temperature. At 25°C, *syp-1^K42E^* animals had almost as few viable progeny as animals lacking an SC altogether [7]. Throughout this work, all experiments were conducted at 20°C, except when analyzing *syp-1^K42E^* animals.

### The N-terminus of SYP-1 is required for complete synapsis

The term ‘synapsis’ has historically been used to indicate both association of the SC with the axis *and* alignment of two axes concomitant with SC assembly. This is indeed the case in wildtype animals, where the SC assembles only between the paired axes, aligning the homologs from end to end. The phenotypes of our mutants (detailed below and summarized in Fig. 2A) show that these two SC activities can be separated. We therefore use the term ‘synapsis’ to describe the association of SC and axes, and ‘zipping’ for SC-mediated alignment of the two axes. In *syp-1*^*Δ8*^ animals zipping is defective and synapsis occurs with unpaired axes. Zipping is also defective in *syp-1^K42E^* animals. At 20°C the SC fails to fully zip the homologs but synapsis is still exclusive to paired axes. However, at 25°C zipping is nearly abolished, and this is accompanied by widespread synapsis with unpaired axes.

**Figure 2.**
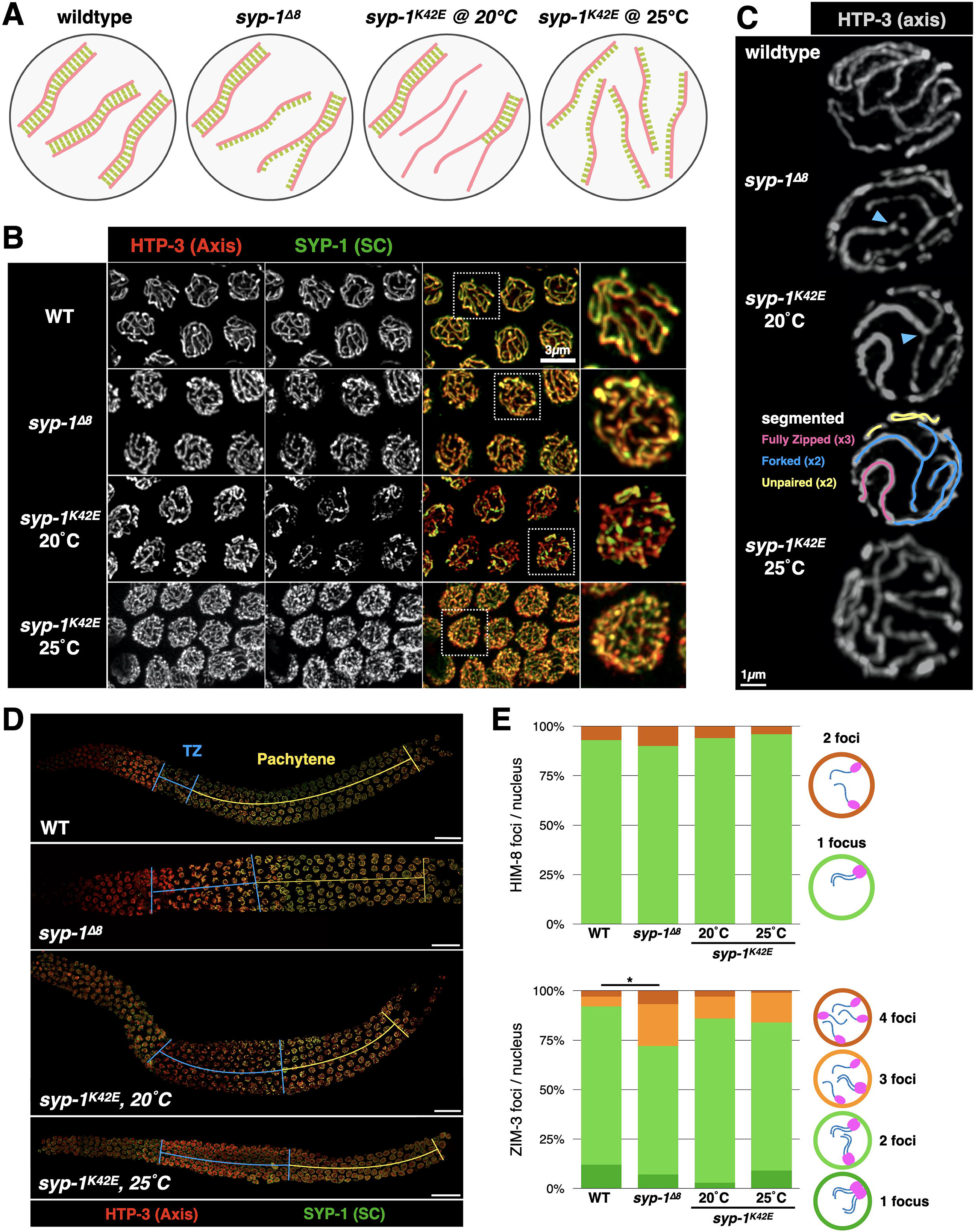
*syp-1* mutants display distinct defects in SC assembly. A) Model of chromosomal organization and SC association in *syp-1* mutants. For simplicity, only three homolog pairs are shown. Salmon, chromosome axis; green, SC; gray; nuclear envelope. B) Pachytene nuclei from wildtype worms and *syp-1* mutants showing synapsis defects. Nuclei are labeled with antibodies against the axis protein HTP-3 (red) and the SC (anti-SYP-1 antibodies; green). Higher magnification images are shown to the right. Notably, the SC fails to fully associate with all chromosomes in *syp-1^K42E^* at 20°C, as revealed by axis signal not associated with SC. Note the numerous SC threads in *syp-1*^Δ*8*^ and *syp-1^K42E^* (25°C), caused by the unpaired chromosomes still associated with the SC. Scale bar = 3μm. C) Partial projections of detergent-extracted pachytene nuclei from wildtype and *syp-1* mutants labeled with anti-HTP-3 (gray). The enhanced resolution allows tracing of the different synapsis outcomes, shown for *syp-1^K42E^* (20°C; skeletonized segments): fully zipped (pink), forked (blue), and unpaired and asynapsed (yellow). Forked chromosomes are indicated with a blue arrowhead. Scale bar = 1μm. D) Immunofluorescence images of axis (anti-HTP-3, red) and SC (anti-SYP-1, green) in the indicated genotypes indicating elongated transition zone in *syp-1* mutants. Nuclei progress in the gonad from left to right as they progress in meiosis. Blue lines correspond to the transition zone based on chromatin morphology and yellow lines designate pachytene. Scale bar = 15μm. See Fig. S1E for quantitation. E) Top, X chromosome pairing shown by localization of the Pairing Center protein HIM-8 in animals of the indicated genotypes. Paired and unpaired X chromosomes are shown as one or two HIM-8 foci, respectively. Models are shown to the right. N>130 nuclei in each condition. Pairwise comparisons to wildtype are not significant (Pearson’s chi-squared test). Bottom, pairing of chromosomes I and IV shown by localization of the Pairing Center protein ZIM-3 in animals of the indicated genotypes. Correctly paired chromosomes I and IV would result in one or two foci (green), whereas three or four foci indicate incorrect pairing (orange). Models are shown to the right. N>90 nuclei in each condition. Significant pairwise comparisons to wildtype are indicated (Pearson’s chi-squared test, were ‘correct’ and ‘incorrect’ configurations were grouped together; p<0.0005).

To analyze synapsis in our mutants, we visualized the SC and the axes in meiotic prophase nuclei using antibodies against SYP-1 and the axis protein HTP-3. In wildtype animals the SC assembles on all chromosomes, thereby colocalizing the SYP-1 and HTP-3 signals (Fig. 2B). In *syp-1^K42E^* (20°C) animals, some chromosomes were not synapsed, as evidenced by HTP-3 staining unaccompanied by SYP-1 staining (Fig. 2B). Higher magnification analysis indicated that in addition to chromosomes that are completely asynapsed or completely zipped, some chromosomes were forked - zipped along part of their length and their ends splayed and asynapsed (Fig. 2C). Given the severe meiotic defects, we were initially surprised to observe complete colocalization of SC and axes in *syp-1^K42E^* (25°C) and in *syp-1*^*Δ8*^ animals (Fig. 2B). Chromosome morphology, however, was distinct from wildtype animals. Chromosomes in *syp-1^K42E^* (25°C) animals were almost all unzipped, resulting in 12 stretches of axis and SC in each nucleus. This phenotype is reminiscent of mutants in the regulatory proteins HAL-2/3 [27, 28], as well as the small C-terminal truncation in *syp-3(me42)* [10]. Chromosome axes in *syp-1*^*Δ8*^ were also all associated with the SC, but closer inspection revealed a combination of unzipped, forked, and fully zipped chromosomes, similar to the conformations observed in *syp-1^K42E^* (20°C; Fig. 2C). Confirming the presence of unzipped but SC-associated chromosomes, the SC localized to the X chromosomes in *syp-1*^*Δ8*^ animals where X chromosomes pairing was prevented by deletion of *him-8* ([32]; Fig. S1D). Notably, we did not observe non-continuous SC stretches or bubbles in any mutant scenario, consistent with cooperative assembly of the SC [33].

Synapsis defects and exposed chromosome axes have been associated with changes in meiotic progression [34]. Consistent with this, we observed a dramatically extended duration of the leptotene-zygotene stage of meiosis (“transition zone”; Fig. 2D and S1E) in *syp-1^K42E^* (20°C) animals. *syp-1*^*Δ8*^ and *syp-1^K42E^* (25°C) animals had a more mildly extended transition zone despite seemingly complete colocalization of axis and SC staining, which might reflect slow association of the SC with the axis or an altered SC ultrastructure that is not evident at the resolution of light microscopy. We also quantified the relative abundance of an axis and an SC component in wildtype and mutant animals and found only minor effects on SC abundance (Fig. S1F), arguing that reduced SC protein levels cannot account for the phenotypes we observe.

We also confirmed that the zipped chromosomes are indeed homologs. We found that chromosomes I, IV and X are mostly correctly paired in *syp-1^K42E^* and *syp-1*^*Δ8*^ animals ([35]; Fig. 2E), indicating that homolog pairing is not dramatically impacted. Furthermore, since zipping by the SC is an essential precondition for crossover formation in worms [7], the presence of some linked chromosomes prior to the meiotic divisions (Fig. 1E) suggests that the SC was assembling between homologs, and that heterologous synapsis is rare.

Our analysis demonstrated that despite their proximity on the N-terminus of SYP-1, our mutants cause distinct chromosomal phenotypes. Underlying these phenotypes are violations of two principles that govern SC assembly: loading only between two axes, and spreading along the entire length of the chromosome. Association with only paired axes is disrupted in *syp-1*^*Δ8*^ and *syp-1^K42E^* (25°C) animals; full zipping to align chromosomes end-to-end is disrupted in *syp-1*^*Δ8*^ and *syp-1^K42E^* (20°C) animals.

### The N-terminus of SYP-1 is crucial for intra-SC interactions

We next wanted to understand how alterations in the N-terminus of SYP-1 bring about such dramatic chromosome-scale effects. Since the SC has proved challenging to reconstitute *in vitro*, we used several orthogonal approaches to measure the biophysical properties of the SC in our mutants. In many organisms, SC material can form organized ‘droplets’ called polycomplexes that lack axes or chromatin, and that appear as stacked SC [3, 18]. In worms, polycomplexes arise when SC material cannot load onto chromosomes due to elimination of the axis protein HTP-3 [36]. In *htp-3* animals there is generally one spherical polycomplex per nucleus throughout meiotic prophase, until late pachytene when the number of polycomplexes per nucleus increases [18]. *syp-1* mutations alter this dynamic. Polycomplexes form later in *htp-3 syp-1^K42E^* (20°C) animals, and at 25°C most worms do not form polycomplexes and instead exhibit diffuse SC material in the nucleoplasm, while a minority forms a few polycomplexes in late pachytene (Fig. 3A-B and S2A). In contrast, *htp-3 syp-1*^*Δ8*^ polycomplexes appear with similar dynamics to those in *htp-3* animals, but are more numerous and are non-spherical.

**Figure 3.**
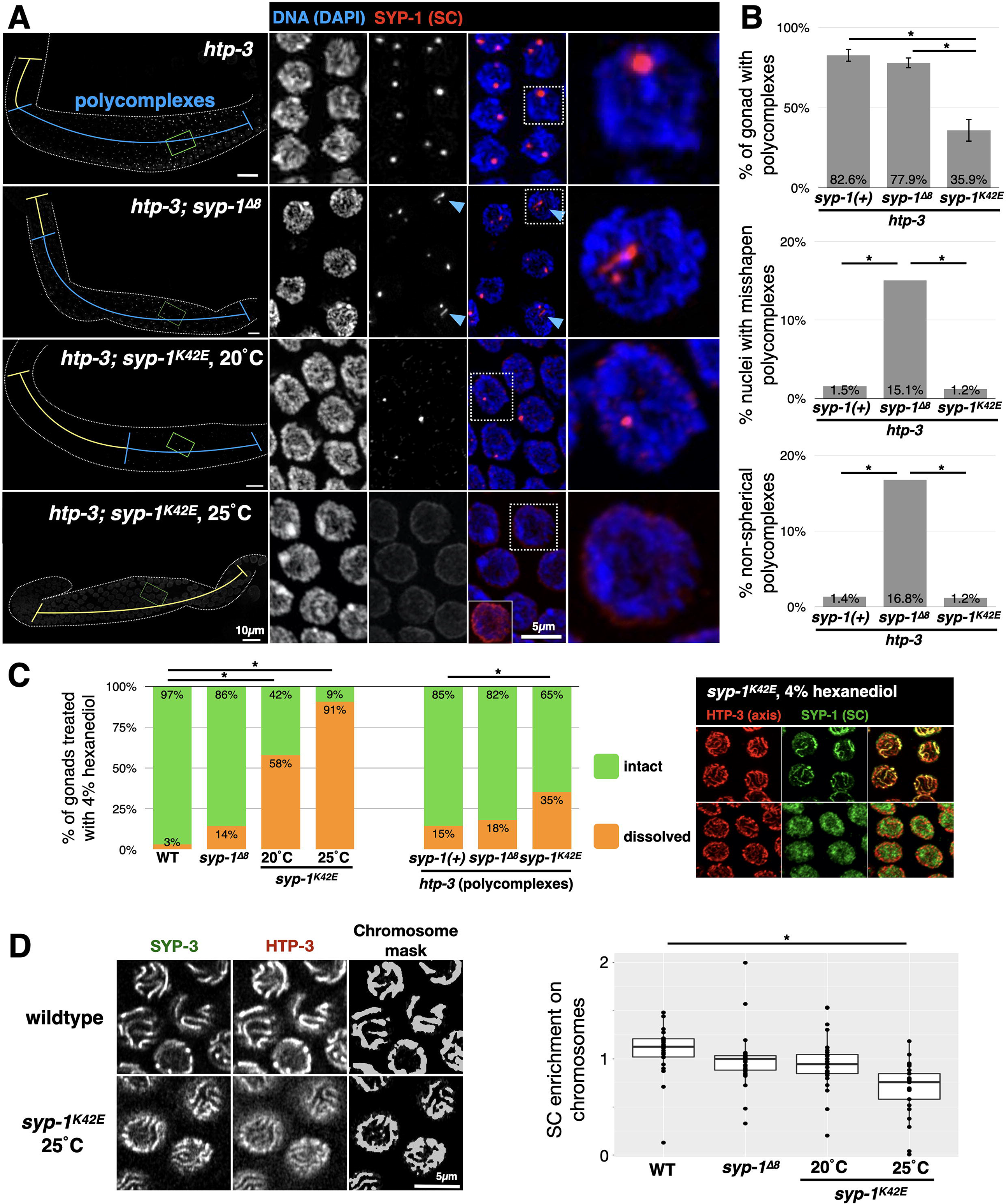
The N-terminus of SYP-1 plays a role in SC-SC interactions. A) Polycomplex morphology and dynamics are altered in *htp-3 syp-1^K42E^* and *htp-3 syp-1*^Δ*8*^ animals. Immunofluorescence images of gonads stained with anti-SYP-1 antibodies. The region of the gonad containing nuclei with polycomplexes is marked with a blue line, whereas the yellow line designates meiotic nuclei without polycomplexes. Scale bar = 10 μm. Middle, pachytene nuclei, with a magnified nucleus shown to the right. SYP-1 polycomplexes are red and DAPI (DNA) is shown in blue. Misshapen polycomplexes (light blue arrows) occur in *htp-3 syp-1*^Δ*8*^ animals. *htp-3 syp-1^K42E^* (25°C) harbor no polycomplexes; bottom right, over-exposed nucleus showing only diffuse SC staining. Scale bar = 5 μm. B) Quantitation of polycomplexes in *syp-1* mutants. Top, the portion of the gonad harboring polycomplexes. Error bars indicate standard deviations. N>4 gonads in each condition. Significant pairwise comparisons are indicated (Student’s t-test, p<0.0005). Middle, percentage of nuclei containing misshapen polycomplexes, scored manually. N>160 nuclei from three different gonads for each genotype. Significant pairwise comparisons are indicated (Pearson’s chi-squared test, p<0.0005). Bottom, non-spherical polycomplexes (ratio between the short and long axis of less than 0.5, where 1 represent a perfect sphere). N>80 polycomplexes from at least three gonads for each genotype. Significant pairwise comparisons are indicated (Pearson’s chi-squared test, p<0.0005). See Fig. S2A for additional data. C) Assembled SC and polycomplexes in *syp-1^K42E^* animals are hypersensitive to dissolution by 4% 1,6-hexanediol. Left, percentage of gonads from the indicated genotypes where the SC was dissolved with 4% 1,6-hexanediol. Middle, percentage of gonads from the indicated genotypes where polycomplexes were dissolved with 4% 1,6-hexanediol. N>30 gonads for each condition. Significant pairwise comparisons to wildtype or *syp-1(+)* are indicated (Pearson’s chi-squared test, p<0.05). Right, immunofluorescence images of representative nuclei with intact SC (chromosome associated) and dissolved SC (diffuse nucleoplasmic) in pachytene nuclei from *syp-1^K42E^* (20°C) animals. D) *syp-1* mutants affect the enrichment of the SC on chromosomes versus nucleoplasm. Left, representative nuclei from wildtype and *syp-1^K42E^* (25°C) animals expressing GFP-SYP-3 and HTP-3-wrmScarlet. A chromosome mask was calculated based on HTP-3-wrmScarlet (see Methods for details). Right, the enrichment ratio was calculated by dividing the sum GFP-SYP-3 intensity inside the chromosome mask by the sum intensity outside the mask, and normalizing by the enrichment value for HTP-3-wrmScarlet. The bars indicate the median enrichment, and the boxes indicate the middle 50%. N>30 nuclei, from at least three animals, for each condition. Significant comparisons to wildtype are shown (p<0.0001; Student’s t-test). Similar enrichment ratios were obtained by calculating peak-to-trough ratios along line scans (data not shown).

We also tested the integrity of the SC, both assembled between chromosomes and as polycomplexes, using 1,6-hexanediol, an aliphatic alcohol that dissolves the SC at high concentrations [18]. At low concentrations, 1,6-hexanediol can be used to probe the strength of phase-separating interactions. In wildtype nuclei, the SC rarely dissolves when treated with 4% 1,6-hexanediol (1/31 gonads exhibited fully dissolved SC; Fig. 3C). The SC in *syp-1^K42E^* animals dissolve more easily with 4% 1,6-hexanediol (41/71 gonads), an effect that is exacerbated at 25°C (29/32 gonads). The SC in *syp-1*^*Δ8*^ animals dissolve at a rate much closer to wildtype (10/70 gonads). Polycomplexes dissolution correlated with that of assembled SC, and was significantly higher for *syp-1^K42E^* compared with wildtype polycomplexes (Fig. 3C).

These analyses of SC and polycomplex integrity suggest that the SC in our mutants, especially in *syp-1^K42E^* (25°C) animals, has weakened intra-SC interactions compared to wildtype. We hypothesized that weaker intra-SC interactions will result in less efficient recruitment of SC subunits onto chromosomes. That was indeed the case: the average enrichment of GFP-SYP-3 on chromosomes relative to the nucleoplasm was ~2-fold lower in *syp-1^K42E^* (25°C) compared with wildtype animals, with *syp-1^K42E^* (20°C) and *syp-1*^*Δ8*^ animals exhibiting similar enrichment to wildtype (Fig. 3D).

Taken together, our analysis indicates that while the chromosomal phenotypes of our N-terminal *syp-1* mutations are related, they stem from different effects on the biophysical properties of the SC. *syp-1^K42E^* weakens intra-SC interactions as reflected by poor assembly into polycomplexes (when loading onto axes is abolished), increased 1,6-hexanediol sensitivity, and inefficient recruitment onto chromosomes, all of which are exacerbated by increasing temperature. In contrast, *syp-1*^*Δ8*^ alters the SC in a way that involves only a minor weakening of SC integrity.

### The N-terminus of SYP-1 plays a role in crossover regulation

The SC regulates crossovers not only by aligning homologs and bringing homologous sequences in close proximity, but also in two additional ways: recruiting pro-crossover factors and regulating crossover distribution [4–6, 18, 25, 26, 37]. Both *syp-1* N-terminal mutants remain competent for recruiting pro-crossover factors – progeny viability and the number of DAPI staining bodies (Fig. 1E) suggest that the SC in *syp-1^K42E^* and *syp-1*^*Δ8*^ animals has not entirely lost the capacity to promote crossover formation. Furthermore, the stereotypic localization of the pro-crossover factor ZHP-3 to polycomplexes – indicative of crossover regulation by the SC independent of the axes or of repair intermediates – was unaffected ([18]; Fig. S2B).

While recruitment of pro-crossover factors remains intact, we wanted to assess whether the number and distribution of crossovers is affected in *syp-1* mutants. We counted GFP-COSA-1 foci, which are cytological markers of crossovers [38], and traced them along each chromosome. GFP-COSA-1 foci were more numerous in *syp-1^K42E^* and *syp-1*^*Δ8*^ worms compared to the six foci observed in wildtype animals, one on each of the six homologs (Fig. 4A-B and Fig. S2C). This was particularly striking in the case of *syp-1^K42E^* (25°C) animals, where despite lack of homolog zipping, and hence of actual crossovers, we observed an average of 16.4 GFP-COSA-1 foci per nucleus (Fig. 4A-B and S2C). Based on observations of differentially labeled sister-chromatids, at least some of these foci likely mark inter-sister exchanges [39]. Tracing of GFP-COSA-1 foci in *syp-1*^*Δ8*^ animals indicated that the increased overall foci number is not due to association of multiple foci per chromosome – like in wildtype animals, every chromosome harbor only a single focus – but likely due to association of foci with the SC on both zipped and unzipped chromosomes (Fig. 4A-B and S2C). In *syp-1^K42E^* animals (at both 20°C and 25°C) many chromosomes harbored more than one GFP-COSA-1 focus, explaining the large number of overall foci (Fig. 4A-B and S2C). To exclude the possibility that a limited number, or altered distribution, of meiotic double-stranded breaks is shaping the crossover landscape in our mutants, we introduced as many as 14 additional breaks per chromosome by X-ray irradiation (Fig. S2D). We observed only minor increases in the number of GFP-COSA-1 foci (Fig. 4C), suggesting that perturbed crossover distribution in our mutants reflects altered regulation by the SC.

**Figure 4.**
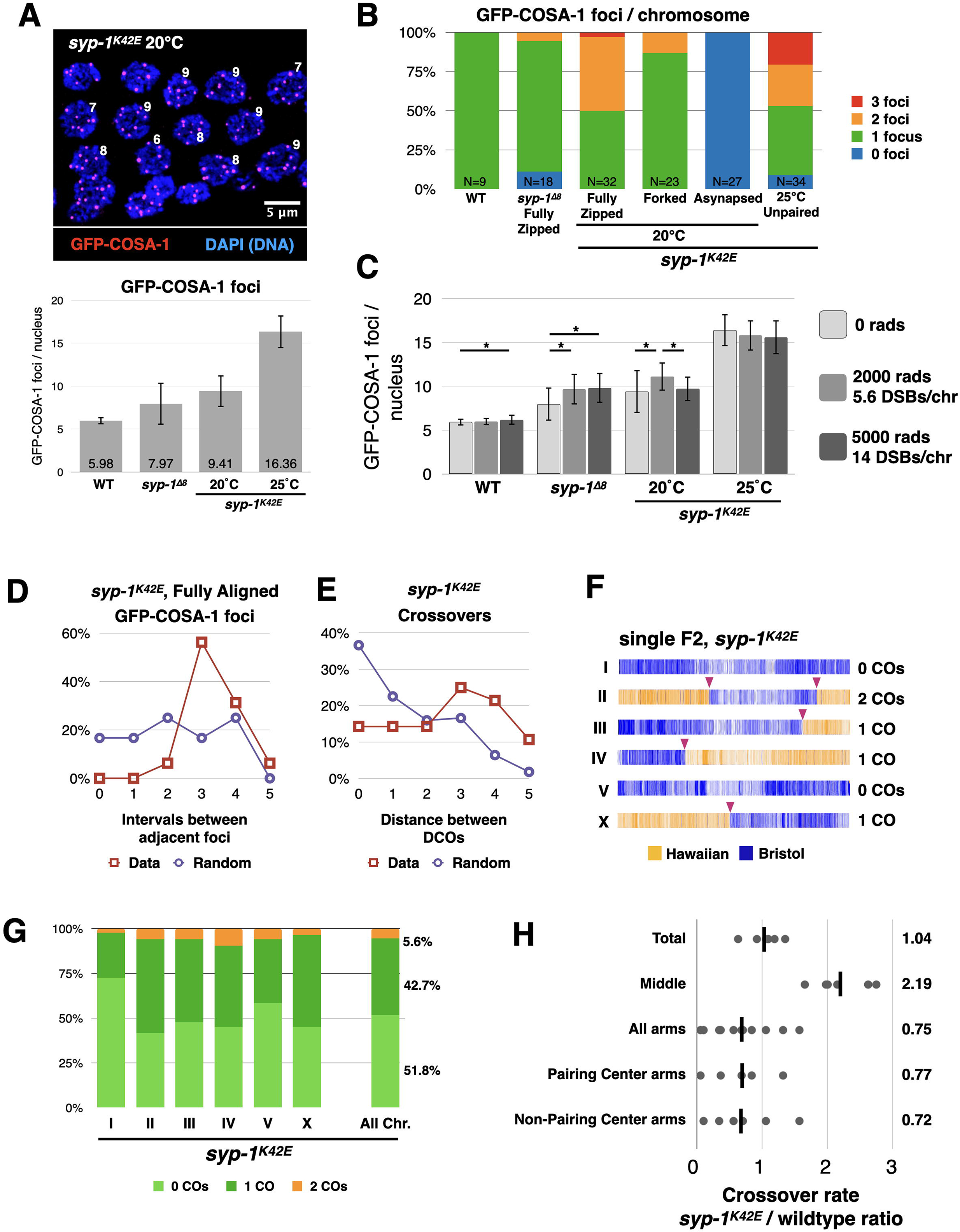
The N-terminus of SYP-1 regulates the number and distribution of crossovers. A) GFP-COSA-1 foci are more numerous in *syp-1* mutants. Top, immunofluorescence images of late pachytene *syp-1^K42E^* nuclei showing the number of GFP-COSA-1 foci per nucleus (red). Scale bar= 5μm. Bottom, the average number of GFP-COSA-1 foci per nucleus in the indicated genotypes. Error bars indicate standard deviation. N>30 nuclei for all conditions. All pairwise comparisons are significant (Student’s t-test, p<0.005). B) The number of GFP-COSA-1 foci per chromosome traced in 3D in the indicated genotypes. Chromosomes in *syp-1^K42E^* animals are shown by the type of chromosome morphology (fully zipped, forked and asynapsed). N values are shown at the bottom of each bar. See examples of traced chromosomes in Fig. S3A. C) The crossover landscape is not limited by the number of double-stranded breaks. GFP-COSA-1 foci in animals of the indicated genotypes after being exposed to varying amounts of X-ray irradiation (2000 and 5000 Rads), creating 5.6 and 14 extra breaks per chromosome, respectively. Error bars indicate standard deviation. N>20 nuclei for all conditions. Significant intra-genotype comparisons are shown (Student’s t-test, p<0.05). D) Intact but reduced interference between crossover precursors. The position of GFP-COSA-1 foci along completely zipped chromosomes in *syp-1^K42E^* animals was divided into six equal bins. In chromosomes that harbor two foci, the foci are more distant from each other (maroon) than would be expected by chance (‘Random’, purple). N=15 chromosomes. E) Interference between genetic crossovers is reduced but intact. In *syp-1^K42E^* chromosome that harbor two crossovers (DCOs), crossovers are more distant from each other (maroon) than would be expected by chance (‘Random’, purple). The chromosomes were divided into 6 equal-sized intervals. N=28 double crossovers. F) SNP profile of a single traceable meiosis (single cross-progeny) showing chromosomes with single, double, and no crossovers (maroon arrows) in *syp-1^K42E^* animals. Each vertical line depicts a unique SNP (Hawaiian, yellow; Bristol, blue), while each horizontal bar represents a single chromosome. G) Prevalence of crossovers (COs) for each chromosome and for all combined chromosomes in *syp-1^K42E^* animals reveals the presence of double crossovers. Crossovers were mapped by sequencing 84 cross-progeny of parental *syp-1^K42E^* animals differing in thousands of SNPs. See Fig. S3C for all mapped crossovers. H) Crossovers in *syp-1^K42E^* animals are shifted towards the middle of the chromosome, but not toward the Pairing Center end. Crossover rate (calculated in centi-Morgan [cM] per Mb) was calculated for each arm and middle of all chromosomes, as defined by [41]. Ratio between the values measured for *syp-1^K42E^* animals and those measured for wildtype animals [41] are shown, with each data point representing a chromosome or a chromosome arm. Note the more than 2-fold higher crossover rate in the middle of the chromosome, and the lower, and much more varied rate on the arms, with no enrichment on the arm harboring the Pairing Center.

The presence of multiple GFP-COSA-1 foci along individual chromosomes in *syp-1^K42E^* animals prompted us to measure crossover interference, which in *C. elegans* acts to ensure a single crossover (or GFP-COSA-1 focus) per homolog pair [4, 38]. GFP-COSA-1 foci in *syp-1^K42E^* animals are farther apart than would be expected if they were randomly distributed (Fig. 4D and S3A), suggesting interference is still intact but acts on a shorter distance than in wildtype. To ensure that crossover interference is indeed affected in *syp-1^K42E^* worms, we identified the location of all crossovers genetically, by crossing *syp-1^K42E^* worms that differ in thousands of SNPs and sequencing the genomes of their progeny (Fig. 4F-G and S3B-C). In *syp-1^K42E^* animals 5.6% of chromosomes harbored double crossovers, higher than the 0% reported for wildtype animals [40]. The lower rate of genetic *versus* cytological double crossovers could be explained by the independent engagement of the four sisters, entailing that only one quarter of cytological double crossovers would manifest genetically. The median distance between double crossovers (14.14 Mb) was much greater than expected if the two crossovers were distributed randomly (3.17 Mb), indicating reduced but active crossover interference, and corroborating our cytological data (Fig. 4D-E).

Our sequencing data revealed additional features of the crossover landscape in *syp-1^K42E^* animals. First, chromosomes formed crossovers at variable rates, ranging from 27% for Chromosome I to 58% for Chromosome II (Fig. 4G), which might reflect variable likelihood of zipping driven by different efficiencies of synapsis initiation on different chromosomes [2, 33, 35]. Second, while crossovers are generally suppressed in the middle of *C. elegans* chromosomes relative to the arms [41], crossovers in *syp-1^K42E^* animals were skewed toward the middle (Fig. 4H). A similar effect was reported in mutants that, like *syp-1^K42E^*, fail to synapse all their chromosomes [42]. Finally, hypothesizing that forked chromosomes reflect zipping that fails to extend to the end of the chromosome, we tested whether crossovers are shifted toward the Pairing Centers, where SC assembly initiates [2, 43]. Surprisingly, we did not observe a shift, but rather idiosyncratic effects on each arm (Fig. 4H). We envisage that synapsis in *syp-1^K42E^* animals initiates at the Pairing Centers and that although forks are relatively stable over short timescales (1hr; Fig. S3D and data not shown), slower SC dynamics over the >24hrs nuclei spend in pachytene shift the zipped regions away from the Pairing Centers. Implicit in this idea is that the local chromosomal environment stabilizes the SC on some regions more than others, thereby facilitating crossover formation in those regions [22].

Altogether, our cytological and genetic analyses demonstrated that the N-terminus of SYP-1 plays a role in crossover regulation beyond zipping homologs together. The SC in both *syp-1*^*Δ8*^ and *syp-1^K42E^* animals can successfully recruit pro-crossover factors, and, when the SC localizes between homologs, can also promote crossover formation. However, the SC in *syp-1^K42E^* animals exhibit significantly weaker crossover interference compared with the SC in wildtype and *syp-1*^*Δ8*^ animals.

Our analysis substantiates the recent understanding of the dual roles of the SC in both promoting and limiting crossovers. In addition to bringing homologs together, recruitment of pro-crossover factors can only happen where SC has assembled, and in that way the SC limits crossovers to synapsed chromosomal regions. This idea receives support in this work, where we show that the SC that assembles on unpaired axes in *syp-1^K42E^* (25°C) animals can recruit GFP-COSA-1 (see also [6, 39]). Our work also strengthens the idea that, at least in *C. elegans,* the SC directly mediates crossover interference. Perturbations of the SC - including partial depletion and mutations in *syp-1*, *-4*, *-5* and *-6* ([4, 11, 13] and this work) - weaken crossover interference. The exact signal that spreads along entire chromosomes to mediate interference remains elusive, but diffusion within a liquid SC has been proposed as a mechanism for how signals might be propagated [18]. Our work strengthens this idea: the correlation between weaker crossover interference and more labile SC in *syp-1^K42E^* animals (Fig. 3) suggests that strong self-association of SC subunits is required for the spreading of the SC along chromosomes *and* for the spreading of an inhibitory signal that limits crossover number. Transient or permanent discontinuities along the SC in *syp-1^K42E^* animals would block spreading of the SC when it assembles, and limit the diffusion of an inhibitory signal. Alternatively, the reduced self-association of SC subunits may alter the recruitment or the diffusion of a signaling molecule (“client protein”) that interacts with the SC, thereby altering signaling dynamics. Future high-resolution imaging efforts to reveal the internal organization of the SC in *syp-1^K42E^* animals will help distinguish between these possibilities and shed light on the mechanisms of crossover interference.

### A biophysical model of SC assembly

Based on our analysis of SC mutants, we propose that the ability of the SC to zip chromosomes relies on two fundamental interactions (Fig. 5A). The first interaction holds the two halves of the SC together (and by extension, the two axes and two homologs). We refer to it as a zipping interaction and propose that it arises from a high energetic cost of exposed surfaces in the middle of the SC. The second is a stacking interaction that allows the SC to recruit new subunits and spread along chromosomes and likely also drives the phase separation of the SC [18]. In this model, zipping interactions are disrupted in *syp-1*^*Δ8*^ animals. As a result, the SC in *syp-1*^*Δ8*^ animals spread along the entire length of all chromosomes, but the chromosomes are not held together strongly and hence fork. The non-spherical polycomplexes assembled in *htp-3 syp-1*^*Δ8*^ animals might also be explained by dominant stacking *versus* zipping interactions in this mutant (Fig. 3A-B). In contrast, the SC in *syp-1^K42E^* animals are mostly defective in stacking interactions, which abort the zipping of the chromosomes, and lead to forks with asynapsed regions. As temperature increases, the stacking interactions diminish further (Fig. 5B), resulting in high concentrations of free SC subunits, and consequently spurious associations with unpaired axes. The progressively weaker stacking interactions also manifest in the weaker tendency to form polycomplexes, the hypersensitivity to 1,6-hexanediol, and the reduced enrichment of the SC on chromosomes (Fig. 3).

**Figure 5:**
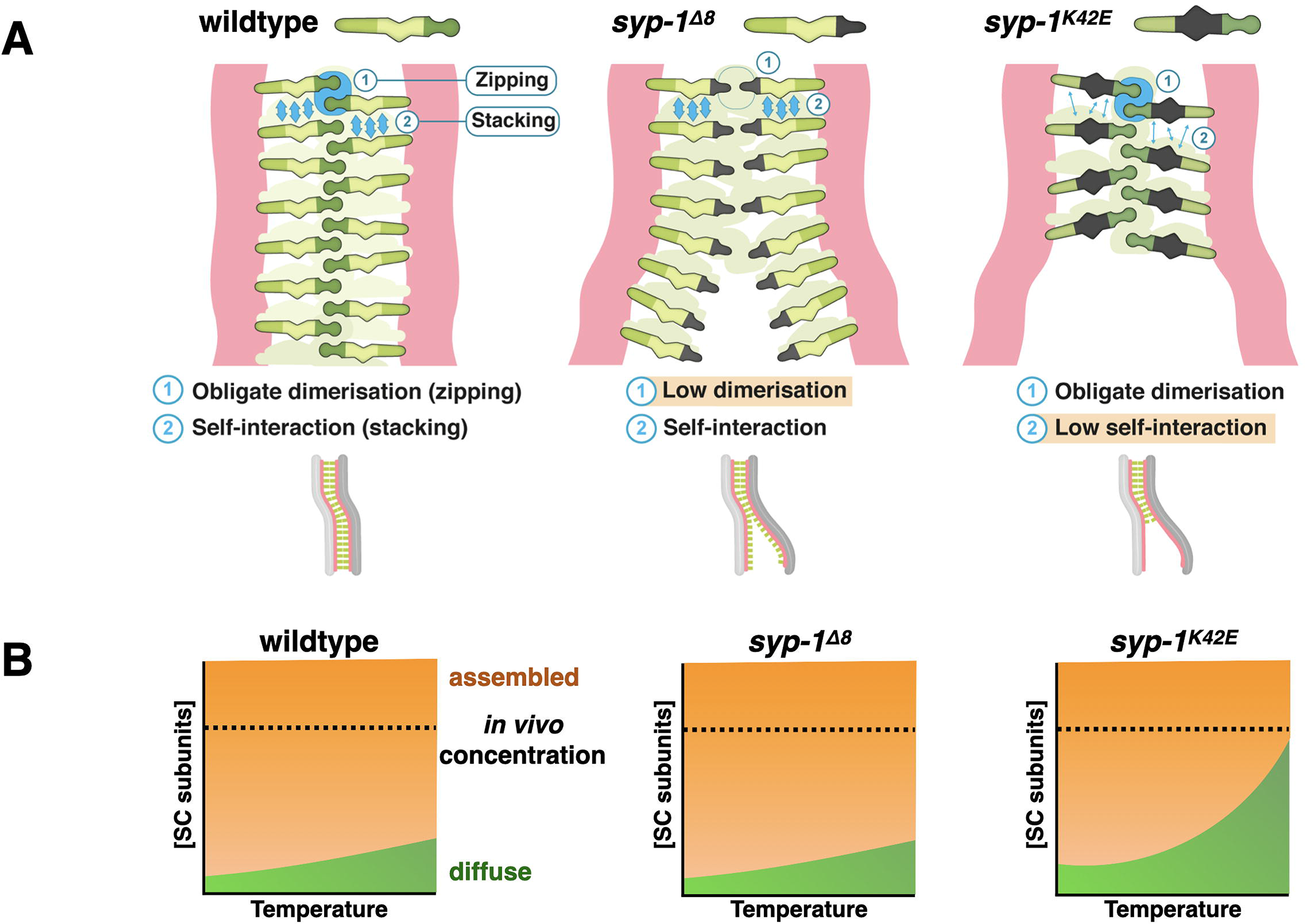
Model. A) Models of the two kinds of interactions allowing the SC to spread. Stacking interactions are shown as horizonal blue arrows, and zipping interactions as vertical arrows. Zipping interactions are weaker in *syp-1*^Δ*8*^ animals leading to inability to hold chromosomes together. Stacking interactions are weaker in *syp-1^K42E^* animals, resulting in incomplete spreading of the SC along the chromosomes, and potential discontinuities within the SC. Bottom, resulting chromosome morphologies in each condition. Salmon, axis; green; SC. B) The weaker stacking interactions of the SC, shown as a phase diagram, where the concentration of the SC subunits is on the y-axis, and temperature is on the x-axis. Orange indicates conditions where SC material assembles, either onto chromosomes or into polycomplexes. In *syp-1^K42E^* animals the phase diagram is shifted so that the SC fails assemble at 25°C with wildtype concentration of SC subunits. The high concentration of free SC subunits in those conditions results in spurious associations with unpaired axes.

What are the molecular underpinnings of these interactions? The location of the SYP-1 N-terminus in the middle of the SC, and the role of a charged residue revealed by the *syp-1^K42E^* mutation, points to a potential charge-charge interface that mediates zipping and disfavors association with unpaired axes. This interface likely overlaps with the amino acids deleted in *syp-1*^*Δ8*^. Stacking of SC subunits is likely to underlie phase separation, which can be driven by weak multivalent interactions [44]. For the SC, several non-mutually exclusive mechanisms have been proposed: weak hydrophobic interactions [18], interactions between the predicted coiled-coils, and interactions between charge-interacting elements (CIEs) [12]. The nature and location of the *syp-1^K42E^* mutation – a charge-altering substitution outside the predicted coiled-coil – argues that hydrophobic interactions or the predicted coiled-coils are not solely responsible for phase separation. In contrast, *syp-1^K42E^* alters an existing CIE, supporting a role for CIEs in driving phase separation of the SC [12, 45]. Future biochemical characterization of protein-protein interactions in wildtype and mutant SC are likely to illuminate the molecular interactions that underlie SC organization.

## Materials and Methods

### Worm strains and transgenes

All strains were grown at 20°C and cultured using standard methods unless stated otherwise [46]. The library of *syp-1* mutants, as well as *syp-1^K42E^* and *syp-1*^*Δ8*^ in the Hawaiian background, were constructed using CRISPR/Cas9 RNP injection with a gRNA and repair template as described below (Fig. S1B-C). Mutants were then crossed out, and linkage between the mutations and the meiotic phenotype was verified. The temperature-sensitive *syp-1^K42E^* was normally maintained at 15-20°C, and grown to adulthood at the indicated temperature to examine its meiotic phenotypes. *syp-1* mutant strains with fluorescently tagged axis and SC components (ROG211 and ROG 250) were maintained as balanced strains, since we detected potential genetic interaction with the EmGFP-SYP-3 transgene, that is itself a hypomorph [2]. *htp-3-wrmScarlet (slc1)* was constructed by CRISPR/Cas9 injection at the C-terminus of *htp-3* using *C. elegans*-optimized Scarlet protein [47]. For a full list of strains used see Table S1. Progeny and male counts were performed by plating an L4 on a new plate each day for three days, then counting progeny and males from each plate.

**Table S1:** Worm strains used in this study.

| Strain | Genotype | Source/Construction |
| --- | --- | --- |
| N2 | Bristol wild type | CGC |
| CB4856 | Hawaiian wild type isolate | CGC |
| CA257 | *him-8 (tm611) IV* | CGC |
| TY5038 | *htp-3 (tm3656) I/hT2 [bli-4 (e937) let-? (q782) qIs48] (I, III)* | CGC |
| ROG120 | *htp-3-wrmScarlet (slc1) syp-3 I; ieSi11[syp-3p::EmeraldGFP::syp-3::syp-3 3'UTR + Cbr-unc-119(+)] II* | Cross between *htp-3-wrmScarlet* and *syp-3 I; ieSi11 II* |
| ROG212 | *mels8 [pie-1p::GFP::cosa-1 + unc-119(+)] II; pSUN-1::TIR-1::mRuby him-8(tm611) spo-11-aid (slc3) IV* | [39] |
| ROG198 | *syp-1<sup>K42E</sup> (slc11) V* | Created by CRISPR/Cas9 microinjections |
| ROG199 | *syp-1<sup>Δ8</sup> (slc12) V* | Created by CRISPR/Cas9 microinjections |
| ROG200 | *htp-3 (tm3656) I; syp-1<sup>K42E</sup> (slc11) V* | Cross between TY5038 and ROG 198 |
| ROG201 | *htp-3 (tm3656) I; syp-1<sup>Δ8</sup> (slc12) V* | Cross between TY5038 and ROG 199 |
| AV620 | *mels8 [pie-1p::GFP::cosa-1 + unc-119(+)] II* | CGC |
| ROG202 | *mels8 [pie-1p::GFP::cosa-1 + unc-119(+)] II; syp-1<sup>K42E</sup> (slc11) V* | Cross between AV620 and ROG198 |
| ROG210 | *mels8 [pie-1p::GFP::cosa-1 + unc-119(+)] II; syp-1<sup>Δ8</sup> (slc12) V* | Cross between AV620, and ROG 199 |
| ROG208 | *him-8(tm611) IV; syp-1<sup>Δ8</sup> (slc12) V* | Cross between CA257 and ROG 199 |
| ROG211 | *htp-3-wrmScarlet (slc1) syp-3 (ok758) I; ieSi11 [syp-3p::EmeraldGFP::syp-3::syp-3 3'UTR + Cbr-* | Cross between ROG120 and ROG198 |
|  | *unc-119(+)* II; *syp-1<sup>K42E</sup>* (*slc11*) V; *nT1 [qls51]* (IV;V) |  |
| ROG250 | *htp-3-wrmScarlet* (*slc1*) *syp-3* (*ok758*) I; *ieSi11* [*syp-3p::EmeraldGFP::syp-3::syp-3 3'UTR + Cbr-unc-119(+)*] II; *syp-1<sup>Δ8</sup>* (*slc12*) V; *nT1 [qls51]* (IV;V) | Cross between ROG 120 and ROG199 |
| ROG255 | *syp-1<sup>Δ8</sup>* (*slc12*) V (Hawaiian) | Microinjections into CB4856 |
| ROG256 | *syp-1<sup>K42E</sup>* (*slc11*) V (Hawaiian) | Microinjections into CB4856 |

### CRISPR/Cas9 injections

L4 worms were grown for 24 hours at 20°C prior to injection. Injection mix was made in the following way: 1 μL *dpy-10* sgRNA (50 μM), 4 μL tRNA (200 μM), 3 μL guide RNA (200 μM) were incubated for 5 minutes at 95°C, then cooled to room temperature. 3.5 μL of the RNA solution was mixed with 3.5μL DNA template (1.5 μg/ml), 0.5μL Cas9 protein (IDT), and 0.5μL of DPY-10 template (200 μM). Dpy and Rol F1 progeny (positive for the co-injection mutation at *dpy-10*) were singled into plates, and their progeny analyzed.

### Immunofluorescence staining and mounting

Gonads were dissected and stained as in [48]. Briefly, 24-h post-L4 adults in egg buffer (25 mM Hepes, pH 7.4, 118 mM NaCl, 48 mM KCl, 2 mM EDTA, 5 mM EGTA, 0.1% Tween-20, and 15mM NaN_3_) and fixed in 1% formaldehyde before freeze cracking on dry ice. Dissected germlines were further fixed in methanol at −20°C for 1 min and rehydrated with PBST. Samples were then blocked with blocking reagent (Roche 01585762001) overnight at 4°C and incubated with primary antibodies overnight at 4°C. Primary antibodies were used at the following concentrations: anti-GFP (mouse, 1:1000; Sigma F1804), anti-SYP-1 (goat, 1:500; [34]), anti-HTP-3 (guinea pig, 1:500; [16]), anti-HIM-8 (rat, 1:500; [32]), anti-ZHP-3 (rabbit, 1:10,000; Abby Dernburg), anti-ZIM-3 (rabbit, 1:250; [35]), anti-H3K36me3 (rabbit, 1:1000; Abcam), anti-DAO-5 (mouse monoclonal, 1:50; DSHB), GFP-booster-Atto488 nanobody (1:200; ChromoTek, Gba488-100). Appropriate secondary antibodies labeled with Alexa Fluor 488, Cy3, Cy5, or Alexa 647 were used at 1:500 dilution (Jackson Immunoresearch). Slides were mounted using fresh mounting media (450 μL NPG-glycerol, 35 μL 2M Tris, and 15 μL water). 1,6-hexanediol treatments were performed as previously describe [18], and each gonad was scored as ‘dissolved’ if SC structures were indiscernible after treatment. The localization of the SC in *syp-1* mutants was also verified by using antibodies against SYP-2 (data not shown).

Detergent extraction (Fig. 2C and S2C) was performed as described in [5]. Briefly, gonads were dissected in 30μl dissection buffer on an ethanol-washed 22×60mm coverslip. After dissection, dissection buffer was removed by pipetting leaving ~5ul on each slide. 50μl of spreading solution was added and gonads were gently spread across the coverslip using a pipet tip. Coverslips were left to dry at room temperature for 1 hour. Then, coverslips were moved to a 37°C heat block for 2 hours. Coverslips were washing in methanol for 20 min at −20°C and rehydrated by washing 3 times for 5 minutes in PBS-T, and immunostained as detailed above. <u>Dissection buffer</u>: 0.1% v/v Tween-20, 0.01% Tetramisole, v/v 85% HBSS. Fix: 4% w/v paraformaldehyde and 3.4% w/v sucrose in water. <u>Spreading solution</u> (for one coverslip): 32ul fix, 16ul 1% v/v Lipsol in water, 2ul Sarcosyl solution (1% w/v Sarcosyl in water),

### Genome wide mapping of crossovers

Hawaiian and Bristol strains differ by thousands of SNPs. *syp-1^K42E^* Hawaiian males were crossed to *syp-1^K42E^* Bristol hermaphrodites. F1s were isolated and backcrossed to *syp-1^K42E^* Hawaiian males, and the F2s progeny of this cross were picked onto individual plates and grown until starved. Genomic DNA from the progeny of 84 F2 animals was isolated using Zymo DNA isolation kit, and was sequenced using a paired end Nextera library preparation.

The median depth coverage for each strain was between 5X and 11X where most strains had an 8X median coverage. Picard (a set of Java command line tools for manipulating high-throughput sequencing data) was used in order to align and determine the quality of sequencing reads. The sequencing data was first trimmed to eliminate the library barcode and the mitochondrial DNA reads were thrown out to give a data set of genomic reads without barcodes. GATK (genomics analysis toolkit) was used for variant calling (identifying SNPs) in the germline using a sequenced *Hawaiian* strain with the *syp-1^K42E^* mutation and the most up to date *C. elegans* Bristol genome as reference genomes. Due to an inferred secondary genome, we manually decided CO points based on how many SNPs in a row and lack of background. For comparisons to the wildtype recombination data (Fig. 4H), we used the CO rates and the designation of that arms and middle of the chromosome defined in [41].

### Irradiation and quantification of breaks

The number of DSBs induced by X-ray irradiation was calculated similarly to the method described in [38]. Briefly, *meIs8 [pie-1p::GFP::cosa-1 + unc-119(+)] II; pSUN-1::TIR-1::mRuby him-8(tm611) spo-11-aid IV* (ROG212) adult hermaphrodites were grown on 1 mM auxin (indole-3-acetic acid, VWR AAA10556-36) and therefore lacked SPO-11. These worms were subjected to increasing dosage of X-ray irradiation using a W-(ISOTOPE) source. 5 hours after irradiation the worms were dissected, and the number of COSA-1 foci, which mark COs, were counted. The results were modeled as in [38], with the exception of assuming a maximum number of 5 foci rather than 6, since the *him-8* mutation prevents X chromosome pairing and synapsis in our background. We solved to minimize the root mean square deviation, yielding a constant of c=0.0028, meaning each 1000 rads of X-ray irradiation caused an average of 2.8 DSBs per homolog pair, or 1.4 DSBs per chromosome.

### Image acquisition and processing

Confocal microscopy images were collected as z-stacks (at 0.2μm intervals) using a 63x NA 1.40 objective on a Zeiss LSM 880 AiryScan microscopy system. Image processing and analysis was conducted using the ZEN software package (Blue 2.1). Partial maximum intensity projections are shown throughout. Foci were counted and confirmed using both maximum intensity projections as well as z-stack images. Transition zone length (Fig. 2D and S1E) was calculated by measuring rows where most nuclei exhibited crescent-shaped chromatin. Meiotic entry was designated by the loading of HTP-3 onto chromosomes, and crescent-shaped was assessed based on chromatin morphology and the location of the nucleolus, marked by DAO-5. The polycomplex zone (Fig. 3B) was calculated by measuring the portion of the gonad from the first nucleus with a polycomplex to the last row of nuclei with a polycomplex. Chromosome tracing, shown in Fig. 2C and used for tracing GFP-COSA-1 to yield the data included in Fig. 4B, 4D, S2C and S3A, was performed on 3D stacks using Imaris 9.2 (Bitplane).

### Fluorescent protein quantification

For overall protein quantification (Fig. S1F), animals expressing GFP-SYP-3 and HTP-3-wrmScarlet (ROG120, ROG211 and ROG250) were picked into 5μL of PEG glycerol (20% (v/v) PEG 20,000 and 20% (v/v) glycerol in PBS) and imaged immediately. The z-stacks were then summed to create a 2D image, and Fiji was used to quantify the individual fluorescent channels. Five pachytene nuclei from each worm were measured and a nuclei sized section of background fluorescence was measured in each gonad and subtracted from the fluorescence measurement. For quantification of chromosome-associated versus nucleoplasmic SC (Fig. 3D), live animals were immobilized and imaged as described in [2], and chromosome masks were calculated by using the ‘Surfaces’ option in Imaris 9.2 (Bitplane). Chromosome-associated fluorescence was calculated by summing fluorescence intensity inside the masked region, and nucleoplasmic fluorescence was calculated by subtracting the chromosome-associated fluorescence from the total nuclear fluorescence. The values obtained for GFP-SYP-3 were normalized to the enrichment values obtained for HTP-3-wrmScarlet. Similar values were obtained by calculating peak-to-trough ratios along line scans taken across single z-planes of pachytene nuclei (data not shown).

### Quantification of polycomplexes sphericity

Sphericity was assessed in Imaris 9.2 (Bitplane). 3D stacks were flattened in the z-dimension to eliminate aberrations caused by the poor resolution, the polycomplexes were automatically segmented using the ‘Surface’ tool, and the “Elipticity (oblate)” measurement was used.

### Alignment of Caenorhabditis species sequences

*Caenorhabditis syp-1* orthologues were identified by using *C. elegans* SYP-1 to query 10 *Caenorhabditis* species genomes using tBLASTn [49] implemented in WormBase or Caenorhabditis.org genome databases. All queries produced a single, high-confidence hit to an annotated gene. The coding sequences of *C. tropicalis* and *C. sinica* required manual editing of one conserved intron-exon boundary. Protein sequences were aligned using ClustalW [50] implemented in Geneious Prime 2019.0.4. Alignments were visualized and colored according to conservation in Jalview [51].

## Authors’ contributions

SGG carried most of the experimental work. LEK performed the detergent extraction and chromosome tracing analysis, and contributed to the phylogenetic analysis. KX discovered that *syp-1^K42E^* animals were temperature-sensitive. SGG and OR conceptualized the project, analyzed the data and wrote the paper, with feedback from all the authors.

## Acknowledgement

We thank members of the Rog lab for critical reading and comments on this manuscript; Abby Dernburg and Yumi Kim for antibodies; Kent Golic for the use of the X-ray source; Yuval Mazor and Yumi Kim for critical reading of this manuscript; Sara Nakielny for comments on the manuscript and editorial work; the Scientific Illustrator Maria Diaz de la Loza for graphical work; and Stuart Cai, Parker Shea, Elom Amematsro and Linda Nikolova for contributions to this project. Worm strains were provided by the CGC, which is funded by NIH Office of Research Infrastructure Programs (P40OD010440). LEK is supported by Developmental Biology Training Grant T32HD007491. This work was supported by R35GM128804 grant from NIGMS and start-up funds from the University of Utah.

---

**Supplementary Figure 1:**
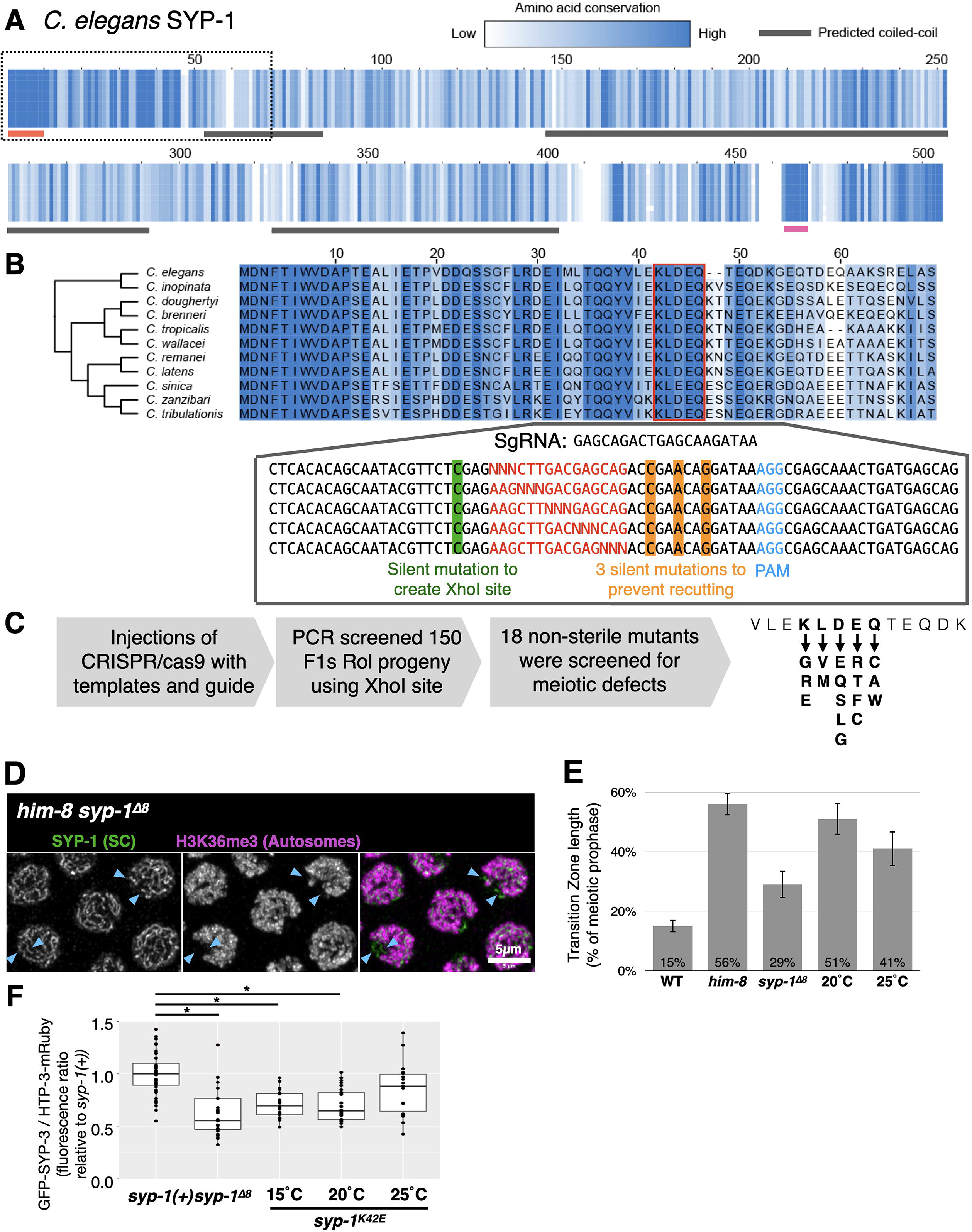
Further characterization of SYP-1 conservation and mutagenesis strategy. A) Sequence alignment of SYP-1 protein sequences from species in the *elegans* group of the *Caenorhabditis* genus with percentages of amino acid conservation (darker blue indicates higher conservation) and predicted coiled-coil domains (dark grey lines below, based on the Paircoil2 program [52]). In addition to the first 13 a.a. in SYP-1, which are likely conserved due to N-acetylation ([31]; red line), and the phosphorylation site near the C-terminus ([29, 30]; pink line), SYP-1^42-46^ is highly conserved. B) The N-terminus of SYP-1 in all of the elegans species group (species tree to the left) with the mutagenized domain highlighted by the red box. Below are the templates used to mutagenize this domain with the guide RNA shown above the PAM sequence in the templates. C) Schematic showing the steps to create the 19 mutants in SYP-1^42-46^ that were screened for meiotic defects. Following injection with a co-injection marker to generate ‘Roller’ animals, heterozygous worms were identified by PCR, restriction enzyme digestion and sequencing, and finally identifying homozygous that were not sterile. D) The SC is associated with the unpaired X chromosomes in pachytene nuclei from *syp-1*^Δ*8*^*him-8* worms. The SC is stained with antibodies against the SC (anti-SYP-1 antibodies; green) and the histone mark H3K36me3 (magenta), which marks chromatin on the autosomes ([53]; the X chromosomes, lacking H3K36me3, are marked with blue arrowheads). Scale bar = 5μm. E) Quantitation of the transition zone length (related to Fig. 2D), calculated as percentages of the gonad relative to the entirety of meiotic prophase. *him-8* animals, where the X chromosome fails to pair and synapse, are shown as a control for highly extended transition zone [32]. Error bars indicate standard deviation. N>4 gonads for each condition. All pairwise comparisons to wildtype are significant (Student’s t-test, p<0.05). F) Ratio of the fluorescence of GFP-SYP-3 and HTP-3-wrmScarlet in the indicated conditions, relative to the average values in wildtype animals. The bars indicate the median fluorescence, and the box indicate the middle 50%. N>20 nuclei for each condition. Significant comparisons to wildtype are shown (Student’s t-test, p<0.05). While the differences are statistically significant, the overall reduction in the protein levels of SC subunits is much smaller that the 60-70% reduction previously shown to affect SC assembly or crossover interference [4, 33].

**Supplementary Figure 2:**
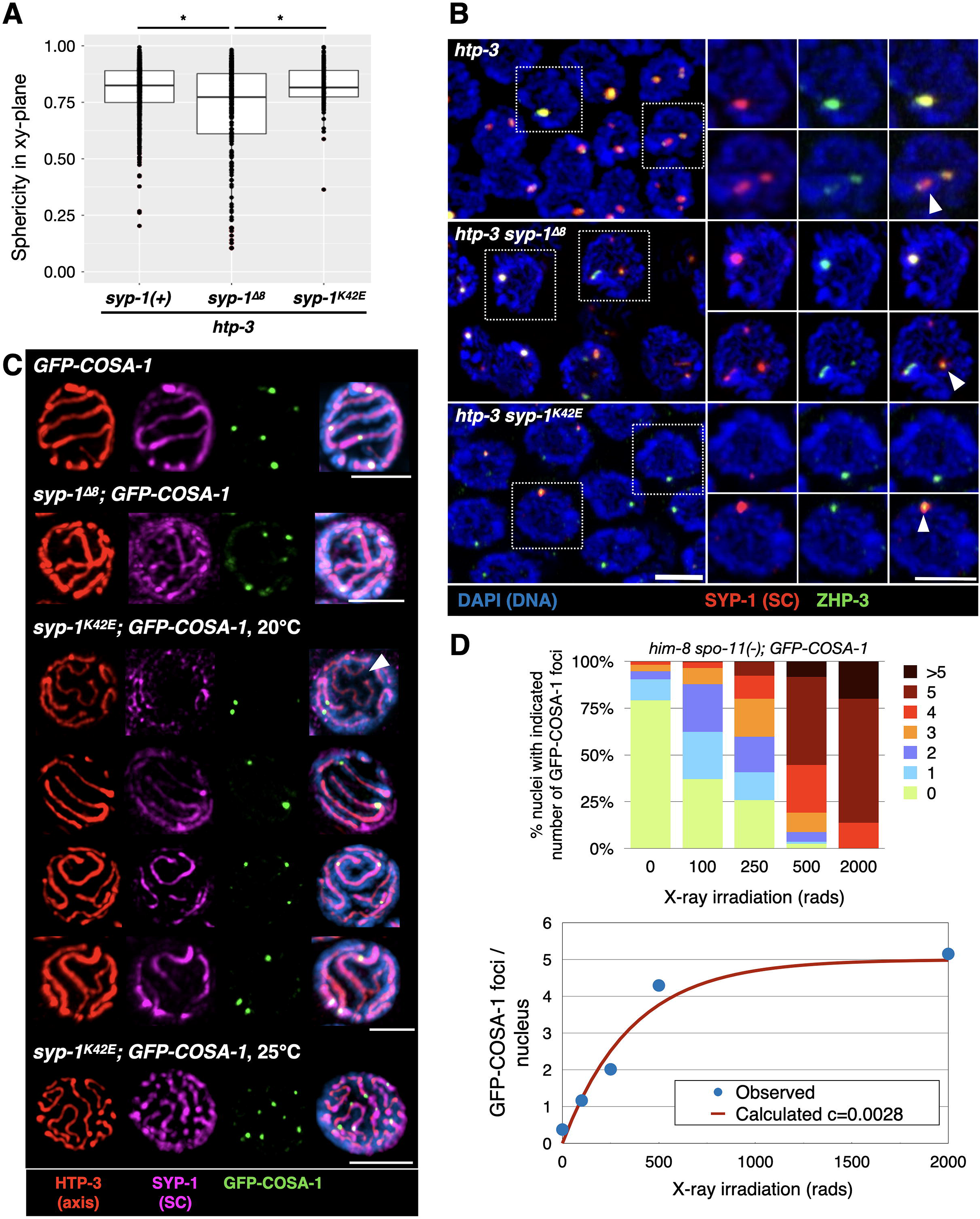
Further characterization of crossovers and polycomplexes in *syp-1* mutants. A) Sphericity measurements of polycomplexes from the indicated genotypes showing that *htp-3 syp-1*^*Δ8*^ has less spherical polycomplexes compared with *syp-1(+)* polycomplexes (related to Fig. 3B). The bars indicate the median sphericity measurement of all polycomplexes from each strain, and the box indicate the middle 50%. N>80 polycomplexes from at least 3 gonads for each genotype. Significant pairwise comparisons are indicated (Student’s t-test, P<0.05). See Methods for more details. B) Polycomplexes in *syp-1* mutants are still able to correctly re-localize ZHP-3. In late pachytene, concomitant with the increasing number of polycomplexes per nucleus, ZHP-3 changes its localization from spreading over entire polycomplexes to small foci abutting polycomplexes [18]. This is also observed in *syp-1* mutants. Immunofluorescence images of the indicated genotypes were stained with anti-SYP-1 antibodies (polycomplexes, red), anti-ZHP-3 antibodies (green), and DAPI (DNA, blue). Right, highlighted nuclei showing both complete overlapping of ZHP-3 and polycomplexes (top), and localization into foci (bottom). Scale bar = 3μm. C) Examples of GFP-COSA-1 localization relative to the SC (related to Fig. 4B). Partial projections of immunofluorescence images of detergent-extracted pachytene nuclei form animals of the indicated genotypes. Nuclei were stained with antibodies against HTP-3 (red), SYP-1 (magenta) and GFP (green; GFP-COSA-1), and with DAPI (DNA; blue). Scale bars = 3 μm. Notably, GFP-COSA-1 foci are associated with the SC in all scenarios. *syp-1^K42E^; GFP-COSA-1* (20°C): Top row, unpaired chromosome that is not associated with the SC (arrowhead) also does not recruit GFP-COSA-1. 2^nd^ row, fully zipped chromosome with a single GFP-COSA-1 focus. 3^rd^ row, a fully zipped chromosome with two foci (arrowhead). Bottom row, a forked chromosome (arrowhead) with a single GFP-COSA-1 focus in the zipped region. Throughout our images we found that GFP-COSA-1 foci were not enriched near forking junctions (only 1/25 junctions was associated with a focus, and only 4 of 25 foci (16%) were <1μm from a junction), arguing against the idea that crossovers locally stabilize the SC. *syp-1^K42E^; GFP-COSA-1* (25°C): Note multiple GFP-COSA-1 foci along single chromosomes. D) Empirical determination of the number of breaks generated per dose of irradiation. Top, animals that cannot make endogenous breaks or pair the X chromosome (*him-8 spo-11(−); GFP-COSA-1*) were subjected to varying doses of X-ray irradiation. In each condition, the number of GFP-COSA-1 foci per nuclei is plotted. As expected, the number plateaus at an average of 5 GFP-COSA-1 foci, since only the 5 autosomes can undergo a crossover. Bottom, the average number of the observed COSA-1 foci are shown as blue dots. The best fitting curve (red line) was calculated as in [38], and was found to result from a conversion factor c=0.0028, meaning each 1 Rad of X-ray irradiation is causing an average of 0.0028 breaks per homolog pair.

**Supplementary Figure 3:**
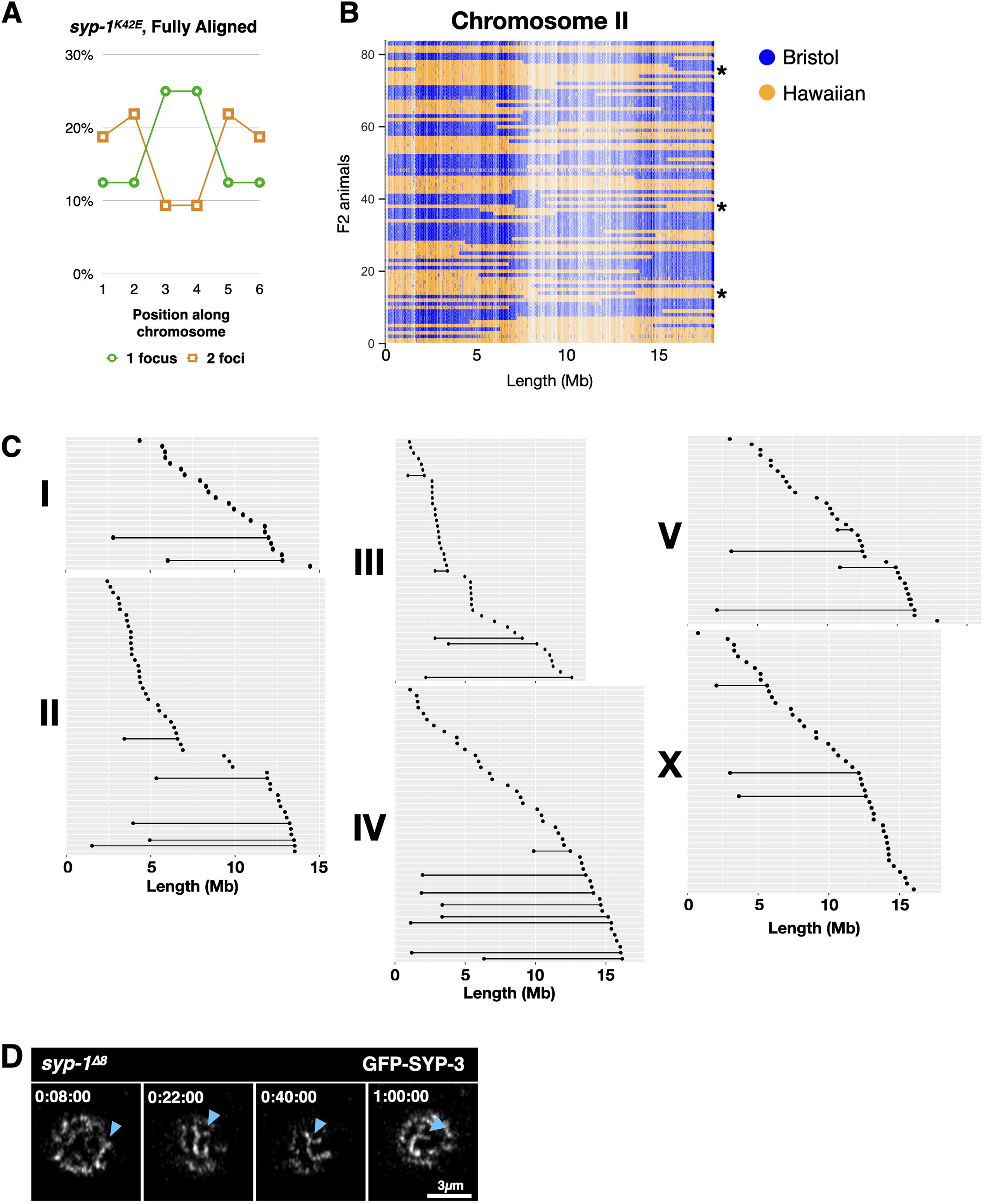
Further characterization of crossovers in syp-1 mutants. A) Intact but reduced interference between crossover precursors. Histograms of the position along completely zipped chromosomes (divided into 6 equal bins) of GFP-COSA-1 foci in *syp-1^K42E^* animals. When 2 foci are present (orange), they tend to be at the opposite ends of the chromosomes. N;15 chromosomes. B) Map of the SNPs along chromosome II in all F2 progeny from an N2 (Bristol, blue) and Hawaiian (gold) cross in the *syp-1^K42E^* background. Note the unique position of every crossover, and the presence of double crossovers in some progeny (several examples noted with asterisks). C) Graphs of all crossovers identifies among the 84 F2 progeny analyzed from the cross described in panel C. Each black dot represents a crossover event, and each horizonal line represents a double-crossover event. The x-axis is the location along each of the six chromosomes. Note the relatively wide spacing of double-crossover events (quantified in Fig. 4E). D) Selected images from a time-lapse series of a live *syp1*^*Δ8*^*GFP-SYP-3* worms. The forked chromosome (white arrowhead) persists throughout the duration of the imaging (1 hour). Times are shown as h:mm:ss. A single z-slice is shown for clarity. Similar results were obtained for *syp1^K42E^ GFP-SYP-3* worms (data not shown). Scale bar = 3μm.

